# All-FIT: Allele-Frequency-based Imputation of Tumor Purity from High-Depth Sequencing Data

**DOI:** 10.1101/625376

**Authors:** Jui Wan Loh, Caitlin Guccione, Frances Di Clemente, Gregory Riedlinger, Shridar Ganesan, Hossein Khiabanian

## Abstract

**Motivation:** Clinical sequencing aims to identify somatic mutations in cancer cells for accurate diagnosis and treatment. However, most widely used clinical assays lack patient-matched control DNA and additional analysis is needed to distinguish somatic and unfiltered germline variants. Such computational analyses require accurate assessment of tumor cell content in individual specimens. Histological estimates often do not corroborate with results from computational methods that are primarily designed for normal-tumor matched data and can be confounded by genomic heterogeneity and presence of sub-clonal mutations.

**Methods:** All-FIT is an iterative weighted least square method to estimate specimen tumor purity based on the allele frequencies of variants detected in high-depth, targeted, clinical sequencing data.

**Results:** Using simulated and clinical data, we demonstrate All-FIT’s accuracy and improved performance against leading computational approaches, highlighting the importance of interpreting purity estimates based on expected biology of tumors.

**Availability and Implementation:** Freely available at http://software.khiabanian-lab.org.

## INTRODUCTION

Clinical sequencing assays aim to identify somatic mutations in cancer cells for accurate diagnosis and treatment of patients through therapeutic targeting of driver alterations and tumor mutational signatures (Garraway, 2013). Although in most scientific settings, patient-matched tumor and germline DNA samples are sequenced for this purpose, most implementations for clinical sequencing lack control DNA data and often only tumor specimens undergo genomic profiling (Frampton, et al., 2013). Moreover, because of recent rulings on qualified coverage for genomic diagnostic assays (United States Food & Drug Administration, 2017), tumor-only sequencing is poised to become one of the most utilized methods for genomic profiling of cancer patients in clinical settings.

Most specimens that are collected in the clinic are formalin-fixed, paraffin-embedded (FFPE), and contain a mixture of tumor cells as well as surrounding non-tumor cells, which include stromal and hematopoietic populations. Hybrid-capture, high-depth sequencing (i.e. >500× depth of coverage) permits identification of genomic alterations with high statistical confidence in measuring variant allele frequencies (VAF), especially compared to whole-genome and whole-exome sequencing methods (Damodaran, et al., 2015; Shaw and Maitra, 2019). Detected variants in tested samples may arise from germline mutations present in all cells, somatic genomic alterations present in all cancer cells, and somatic genomic alterations present in a subset of cancer cells or occasionally in a subpopulation of non-tumor cells (Severson, et al., 2018). The power to detect these small somatic clones depends on sequencing depth and relative abundance of each cell population that harbors them. In the absence of patient-matched control DNA, the Single Nucleotide Polymorphism database (dbSNP) as well as variants detected in healthy individuals within public or private cohorts are often used to identify and remove germline variants (Li, et al., 2017). However, this approach fails to capture rare germline alterations that are specific to each patient, leading to possible misclassification of germline mutations as somatic. Therefore, determining whether detected a mutation is germline or truly somatic as well as resolving variant clonality and loss of heterozygosity (LOH), require additional analyses based on an accurate estimation of specimen’s tumor content or purity. LOH events are shown to be pertinent for assessing treatment efficacy (Brok, et al., 2017; McGranahan, et al., 2017; Pawlyn, et al., 2018; Sade-Feldman, et al., 2017). Yet, without accurate estimates of purity, even the availability of patient-matched germline data may not help resolve evidence of LOH in the tumor, which can occur by deletion of the wild-type copy or by duplication of the mutant allele with the loss of the wild-type (copy-neutral LOH). Moreover, accurate estimates of specimen purity help resolve genomic diversity in both tumor and non-tumor cell populations, which adds to the complexity of interpreting a tumor’s mutational landscape and confounds distinguishing sub-clonal tumor alterations from those in a subpopulation of non-tumor cells (Ptashkin, et al., 2018; Riedlinger, et al., 2019; Severson, et al., 2018). In particular, characterizing clonal mutations, which are present in all cancer cells and are postulated as the best candidates for targeted treatment versus mutations that are sub-clonal and are present only in a subpopulation of cells (Amirouchene-Angelozzi, et al., 2017), is contingent upon correct estimation of the tumor content in sequenced specimen.

Histological approximations of tumor purity are not always available or when they are, they do not provide the required confidence for these analyses. For this reason, various computational algorithms have been developed that utilize patient-matched germline sequencing or process-matched normal control data to infer tumor content of tumor specimens as well as genome-wide ploidy within the tumors (Yadav and De, 2015). A few methods have been developed to simultaneously estimate tumor purity and average ploidy from somatic DNA aberrations; however, the mathematical models used in some of the most utilized methods are optimized for SNP array platforms (Carter, et al., 2012; Van Loo, et al., 2010). Other approaches such as CNAnorm (Gusnanto, et al., 2012) and Control-FREEC (Boeva, et al., 2012) assess copy-number and tumor ploidy of a specimen by correcting its contamination with normal cells, normalizing and scaling the data across tumor genomes. Nevertheless, these approaches operate under the assumption that most tumors are composed of a single clone, which causes the estimates to be inaccurate for heterogenous tumors with multiple subpopulations. Other algorithms such as Pyclone (Roth, et al., 2014) and EXPANDS (Andor, et al., 2014) utilize tumor purity and based on a clustering of somatic mutations with similar cellular prevalence, they predict clonal size and mutations specific to each subpopulation. Regardless of the strengths or weaknesses of these methods, they are primarily designed for normal-tumor pairs, and when applied to high-depth, tumor-only data from targeted genomic regions, they produce results that are confounded by the limited scope of sequencing as well as genomic heterogeneity, aneuploidy, and presence of sub-clonal mutations in cancer cell populations. Therefore, there is a need for strategies that take advantage of the power of tumor-only clinical sequencing assays for high-confidence measurements of VAF and focal copy-number variations (CNV). To this end, we developed All-FIT (Allele-Frequency-based Imputation of Tumor Purity), a weighted least square method that through iterative steps estimates specimen purity and its associated confidence intervals using detected variants’ VAF and CNV. We evaluate All-FIT’s performance using a comprehensive set of simulated datasets and compare its results with those from ABSOLUTE (Carter, et al., 2012), the leading algorithm for estimating purity from VAF and CNV measurements in matched normal-tumor data. Finally, we apply All-FIT to high-depth sequencing of 1,861 specimens from patients with solid tumors and show that histological estimates of purity often do not correspond to observed VAF of detected variants, especially when their biological nature is considered. Specifically, we demonstrate the concordance of our estimates with expected biology of tumors, by focusing on specimens from a wide range of cancer types that harbor known hot-spot mutations in the *TP53* gene as well as colon cancer specimens that carry pathogenic substitutions or indels in the *APC* gene, highlighting the prevalence of LOH that affect these commonly mutated tumor suppressors.

## MATERIALS & METHODS

### All-FIT weighted least square approach of imputing tumor purity

All-FIT considers all detected variants as its input and requires their observed VAF (*f*), their total sequencing depth (*D*), and their loci’s chromosomal copy-number or ploidy (*Y*). We assume the positions of all mutations have chromosomal copy-number of 2 in normal cells. Because the germline-vs-somatic status of the detected variants is yet to be determined, we need to evaluate the likelihood of *Y* somatic and *Y* germline mutational models with their corresponding mutated allele’s copy-number *c*_m_ (1≤*c*_m_≤*Y*) for each variant. For a given purity (*p*), we calculate cancer cell fraction (CCF) for somatic mutations as the ratio of observed VAF divided by the expected VAF. Although CCF for unfiltered germline heterozygous mutations is always equal to 1, CCF for LOH, copy-neutral LOH, and amplification events at the loci of germline mutations can be calculated and generalized as a function of *p* (Figure 1). Intuitively, at correct estimate of *p*, CCFs are equal to one for the variants’ likeliest mutational model. Therefore, we postulate that if detected variants are clonal, the parsimonious estimate of tumor content is the value that optimizes 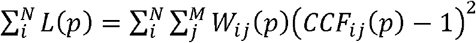; here, *i* and *j* count *N* variants and *M* mutational models, respectively, and *W*_*ij*_ is the Akaike Information Criterion (AIC) weight of mutational model *j* for variant *i*, calculated based on binomially distributed variant depths. To calculate *W*_*ij*_, we follow the approach previously implemented in the LOHGIC algorithm (Khiabanian, et al., 2018), where AIC weights are based on binomial likelihoods of observing a variant with a VAF of *f* and focal ploidy of *Y* at total sequencing depth of *D* across 2*Y* possible mutational models.

**Figure 1.**
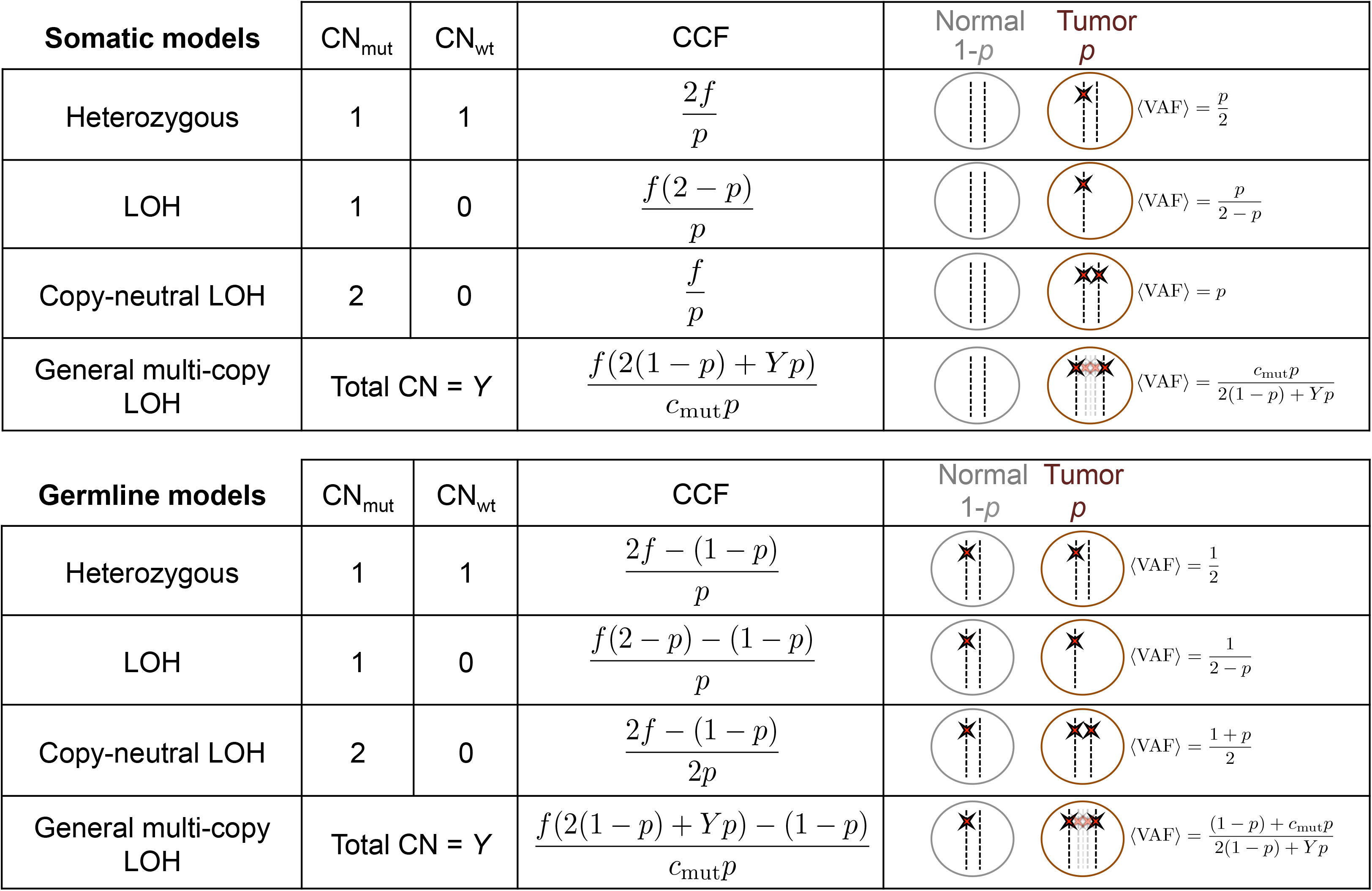
Modeling mutational status and the number of mutated alleles for somatic and germline mutations with their respective expected variant allele frequency (VAF) and cancer cell fraction (CCF). Adapted from Khiabanian *et al.*, 2018.

Since most tumors are heterogeneous in nature and contain both clonal and sub-clonal alterations, we need to evaluate variants’ clonality, which is not possible when *p* is unknown. Therefore, we propose to first use all detected variants to obtain an estimate for purity that can guide the identification and exclusion of unfiltered germline heterozygous and sub-clonal somatic variants. Next, using the remaining putatively clonal events, we optimize *L* to estimate *p* and its confidence intervals (Figure 2a). All-FIT is implemented in Python 3 and is freely available at http://software.khiabanian-lab.org, along with the scripts to generated simulations.

**Figure 2.**
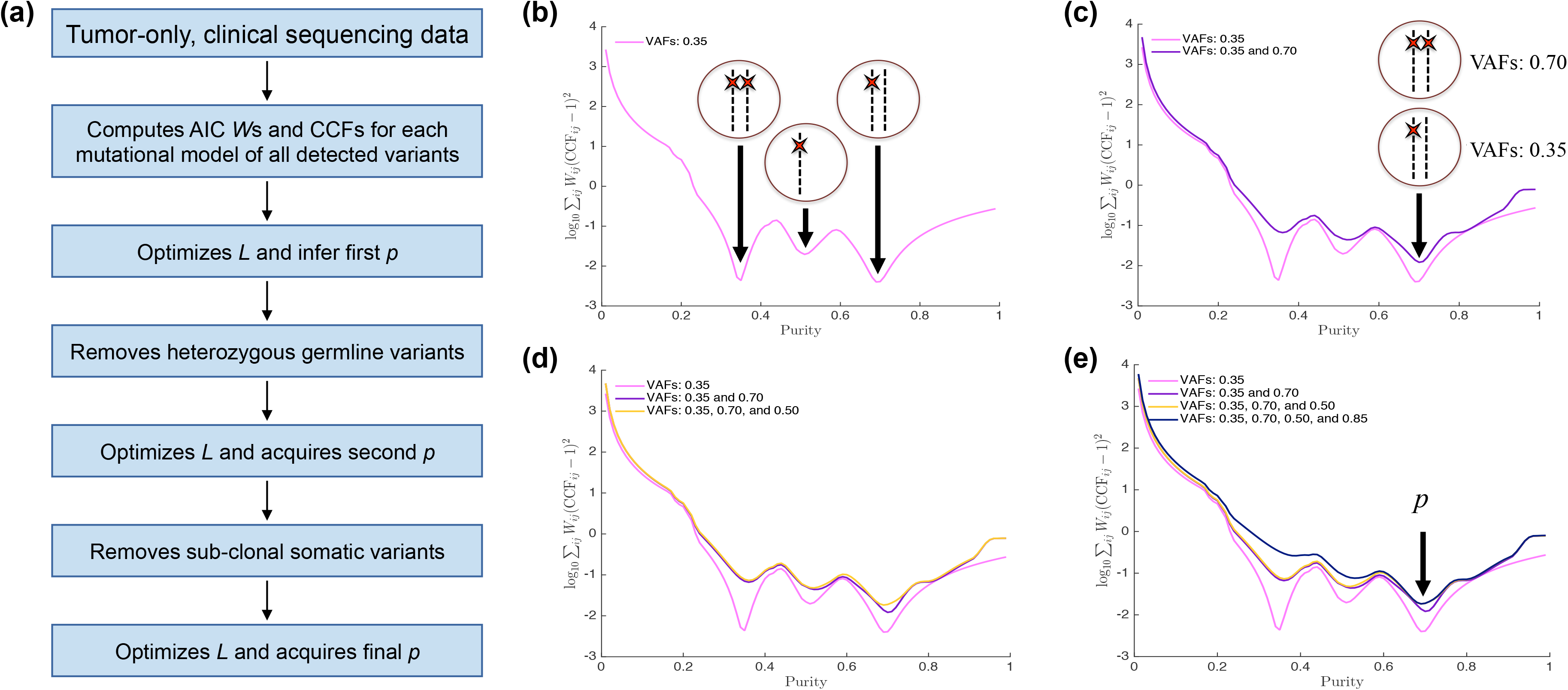
Schematic view of All-FIT’s implementation. (a) All-FIT follows three steps for estimating *p*, first assuming all variants are clonal, then removing germline heterozygous variants, and finally excluding sub-clonal somatic variants. (b) If only a group of variants with observed VAFs of 0.35 exists in a specimen, these variants cannot be distinguished between three different somatic mutational models of heterozygous, under LOH, or under copy-neutral LOH, without knowing specimen’s tumor content. (c) If a second group of variants is detected with observed VAFs of 0.70, *p* = 0.7 can classify variants with observed VAFs of 0.35 and 0.70 as heterozygous and copy-neutral LOH somatic mutations, respectively. (d,e) Detection of additional variants with observed VAFs of 0.50 and 0.85 improves confidence in estimating purity.

### Simulated data

To evaluate All-FIT’s performance, we generated 10,000 simulated sets of variants for known values of *p* (range: 0.1 – 0.9), including a mixture of clonal and sub-clonal mutations with varying ploidy values (dataset 1). The number of variants for each simulated set ranged from 20 to 100 for which we considered equal probability to be under the following eight mutational models: somatic models with heterozygous, LOH, copy-neutral LOH, and high-ploidy alterations, as well as germline models with heterozygous, LOH, copy-neutral LOH, and high-ploidy alterations. These sets included at least one somatic heterozygous mutation; at most 25% of somatic heterozygous mutations were assigned to be sub-clonal. We randomly assigned to each variant a sequencing depth, *d*, ranging from 300× to 1000× (uniformly distributed). Each variant’s observed VAF was randomly generated from a binomial distribution using *d* as the number of trials and expected VAF of variant’s assigned mutational model as the success probability.

To assess the broad utility of All-FIT, we also generated two other datasets with similar conditions. Dataset 2 was enriched with sub-clonal mutations assumed to be somatic heterozygous variants with *c*_m_ = 1. To enrich simulated sets with sub-clonal mutations, we increased the number of somatic heterozygous variants to at least 25% of total number of variants and required at most 67% of them to be sub-clonal. Dataset 3 was enriched with high ploidy mutations (i.e. variants with 3 ≤ *Y* ≤ 8 and *c*_m_ ≥ 1), where we simulated approximately 100 sets for each percentage of high ploidy mutations, ranging from 0% of the variants (absence of high ploidy variants) to 99% (almost all variants in the sample are high ploidy changes).

### Patient data

Patient samples are composed of tumor FFPE specimens submitted for clinical sequencing using the FoundationOne assay (Foundation Medicine, Inc., Cambridge, MA). These samples were sequenced with Illumina Hiseq at >500× on libraries that were enriched with hybrid-capture, using custom bait-sets that target exons and selected introns of 315 genes; after sequencing, these data were analyzed with a process-matched normal control, which is an internally-validated mixture of 10 heterozygous diploid HAPMAP samples, that facilitated normalization of sequence coverage distribution across baited targets. All types of genomic mutations, such as substitutions, small indels, rearrangements, and copy-number changes, were identified for each specimen as previously described (Frampton, et al., 2013). All specimens were reviewed by board-certified pathologists at Foundation Medicine to assess tumor purity. Moreover, empirical Bayesian sampling methods were employed to computationally estimate tumor purity and base ploidy.

Anonymized patient-matched clinical and sequence data were acquired from the Cancer Institute’s Bioinformatic Data Warehouse under IRB protocols of 2012002075, 2017000027, and 20170001364 (Foran, et al., 2017). Data included age at diagnosis, cancer type, clinical history, and deep sequencing data.

### Comparing performance with ABSOLUTE

ABSOLUTE (Carter, et al., 2012) is the most widely used computational method to estimate tumor purity and ploidy from somatic variants, especially in the form of copy-number alterations. It requires an input of CNV and optionally single nucleotide variants. To compare All-FIT’s performance with ABSOLUTE, we distributed simulated variants around the coding genome and provided additional information on chromosomal location to the algorithm. Since ABSOLUTE requires only somatic variants from matched-normal data, we excluded all simulated germline variants. We simulated CNV based on all variants with *Y* not equal to 2 and generated one-exon-sized, 99 base-pair segments to reflect higher local ploidy. (Equal length for CNV segments implied that they contributed equal weight in the analysis). Furthermore, copy-number neutral regions were added into the simulated sets, with number of exons identified in the region as the number of probes within the segments. We used ABSOLUTE’s default parameters and chose estimated tumor purity with the highest likelihood for a model with genome-wide ploidy of two.

## RESULTS

Intuitively, All-FIT imputes *p* by choosing the value that classifies most clonal variants into their respective most likely mutational models. Therefore, it requires variants with various models to break ambiguous inferences to estimate *p* with high statistical confidence (i.e. small confidence interval). For example, if we assume a ploidy of 2 in both normal and tumor cells for a variant with observed VAF of 0.35, the variant can be considered as a somatic mutation that is heterozygous when *p* = 0.70, under LOH when *p* = 0.52, or under copy-neutral LOH when *p* = 0.35. If all variants in a simulated set have observed allele frequencies of about 0.35, it will be impossible to infer a purity estimate that distinguishes mutational models (Figure 2b). If the dataset also includes variants with observed VAF of 0.70, the ambiguity between three possible purity estimates can be broken because at *p* = 0.70, variants with observed VAF of 0.35 are classified as somatic heterozygous mutations while variants with observed VAF of 0.70 are classified as somatic mutations with copy-neutral LOH (Figure 2c-e). Germline heterozygous variants with VAF of ~0.50 play minimal role in breaking the ambiguity between different purity models, as their CCF is always equal to one, while sub-clonal somatic variants can confound this approach as by definition, their CCFs are never equal to one.

All-FIT provides graphical presentations of 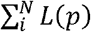 for individual variants (1 to *N*) and presents estimated *p* and its confidence interval at each step, demonstrating how All-FIT imputes *p* often with a higher certainty by removing germline heterozygous and sub-clonal somatic variants. Using the estimated purity, All-FIT calculates the expected VAF from the likeliest mutational model to which each variant belongs, based on an implementation of the LOHGIC algorithm, presenting the concordance between the expected and observed VAFs. It also shows the distribution of CCFs, demonstrating the clonal distribution of variants at the estimated *p*. Ideally, at correct purity, CCF distribution should peak at one with some sub-clonal populations detected at lower fraction, if there are any. Nevertheless, some variants may exist at a fraction that is larger than 1, due to ambiguity in their most likely mutational model (Figure 3).

**Figure 3.**
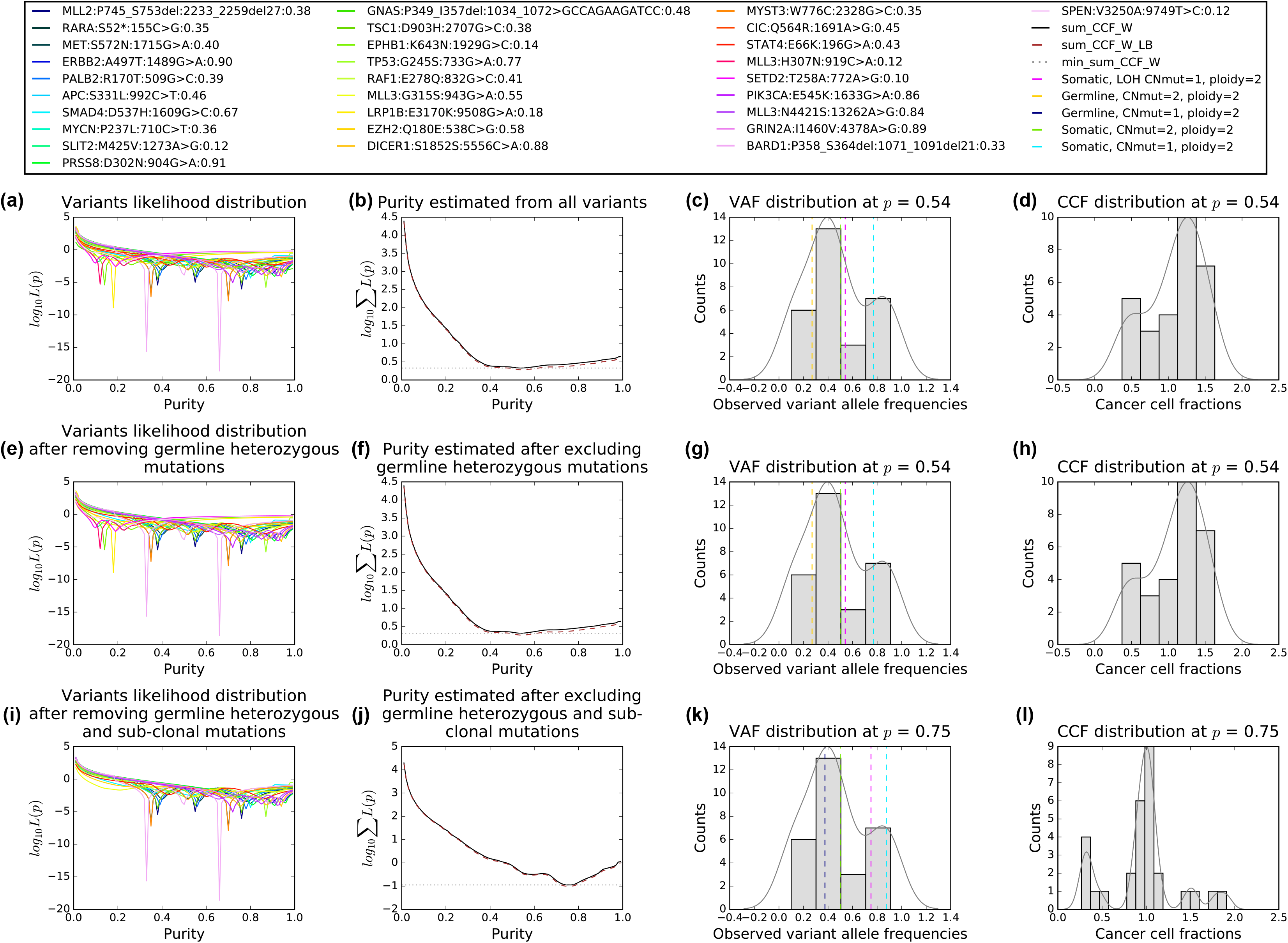
Representation of All-FIT’s results for detected variants. (a) The contribution of each variant to the sum of likelihoods. (b) The sum of likelihoods across all variants, along with its 2σ curve (brown dashed line). The intersections of the 2σ curve and the line tangent to the log of the sum of likelihoods at its minimum (grey dotted line) indicate the confidence interval around the estimated *p*. (c) The distribution of observed allele frequencies. Dashed lines represent the expected allele frequencies of included mutational models for the estimated *p*. (d) The distribution of CCFs for the estimated *p*. (e-h) Results after excluding unfiltered germline heterozygous variants. (i-l) Results after excluding somatic sub-clonal variants.

### Purity estimation in simulated data

We used simulated datasets to assess the accuracy of All-FIT (**Methods**). For dataset 1, comprised of both clonal and sub-clonal mutations with varying ploidy, All-FIT’s purity estimates corroborated with simulated values in 86% of cases with Pearson’s correlation coefficient (*r*) of 0.99 (Figure 4a). All-FIT’s estimates were independent of the total number of mutations in a sample, as the difference between estimated and simulated purities mainly centered around zero especially when there were at least 15 variants per sample. (**Supplementary Figure S1**). When we removed all germline variants from dataset 1 to simulate presence of matched-normal DNA, All-FIT’s accuracy improved to 92%, indicating the robustness of the method by only considering somatic alterations (**Supplementary Figure S2**). In contrast, ABSOLUTE’s estimated *p* agreed with simulated *p* in only 35% of cases with *r* = 0.69 (Figure 4b). It should be noted that assessing All-FIT’s estimated values within their 2σ confidence intervals provided additional predictive power compared to ABSOLUTE, which produces a single purity estimate without a confidence interval.

**Figure 4.**
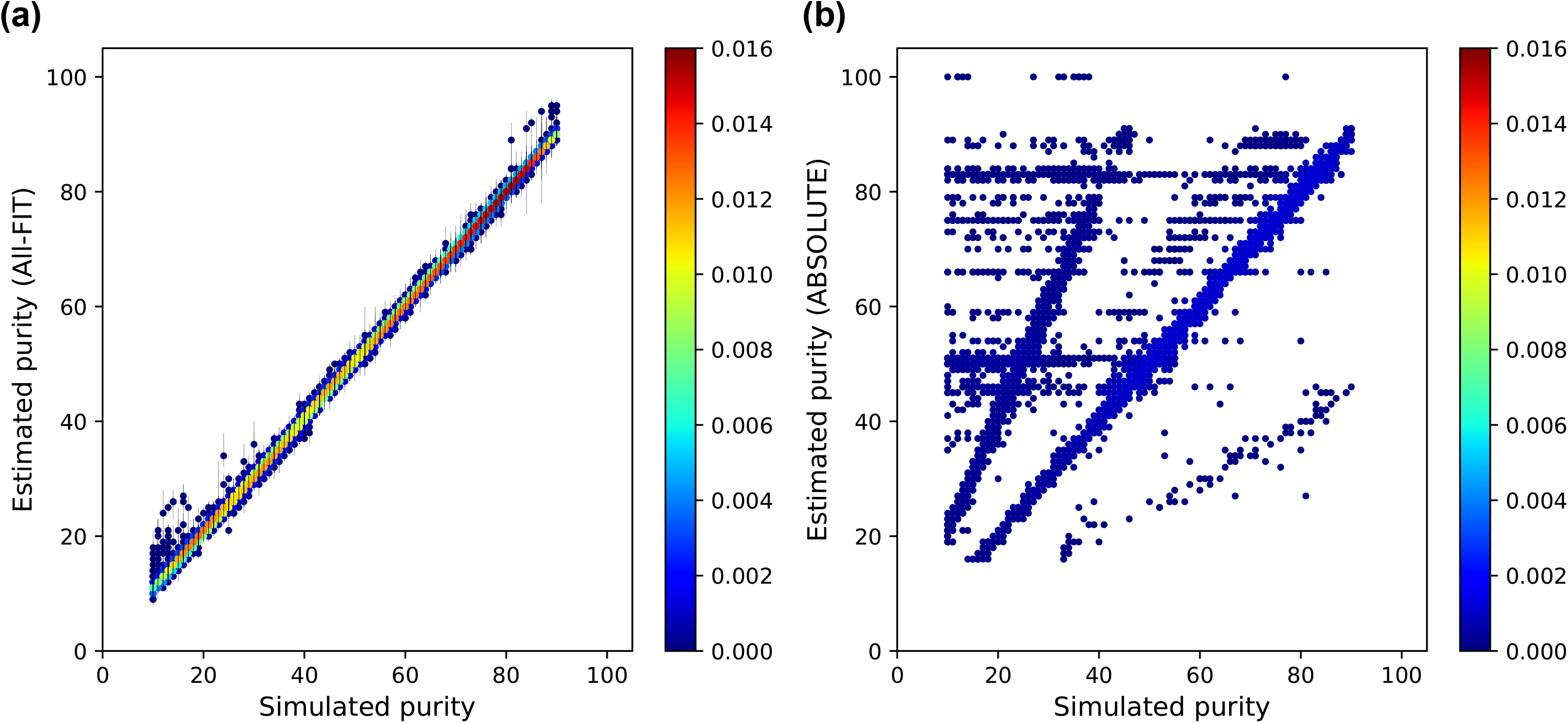
Relationship between simulated purity (ground truth) and estimated purity respectively from (a) All-FIT and (b) ABSOLUTE using dataset 1. The heatmap scale represents the density of data points at each coordinate in 10,000 simulated sets.

Next, we increased genomic heterogeneity in simulated data by varying the number of sub-clonal mutations. We classified the simulated sets into four categories based on the percentage of sub-clonal mutations: <25%, 25-50%, 50-75%, and >75%. All-FIT’s accuracy was not affected by the presence of sub-clonal mutation, when the simulated sets were comprised of <25% of sub-clonal mutations. However, when the percentage of sub-clonal mutations varied from 25% to 75%, All-FIT increasingly misestimated *p* by about one-half, and when more than 75% of the mutations were designated to be sub-clonal, All-FIT estimated purity as one-half of simulated values for most cases. Nonetheless, the correlation coefficient remained higher than 0.76 regardless of the proportion of sub-clonal variants (Figure 5a), and it further improved to 0.84 when we removed germline variants, excluding the 38 simulated sets that had >75% of sub-clonal mutations (**Supplementary Figure S3**). On the other hand, correlation coefficient for ABSOLUTE’s estimates was lower than those of All-FIT, with its best performance of *r* = 0.75 when only <25% of variants were sub-clonal. It was also susceptible to misestimating purity at one-half of the simulated *p*, even with a few sub-clonal variants present (Figure 5b).

**Figure 5.**
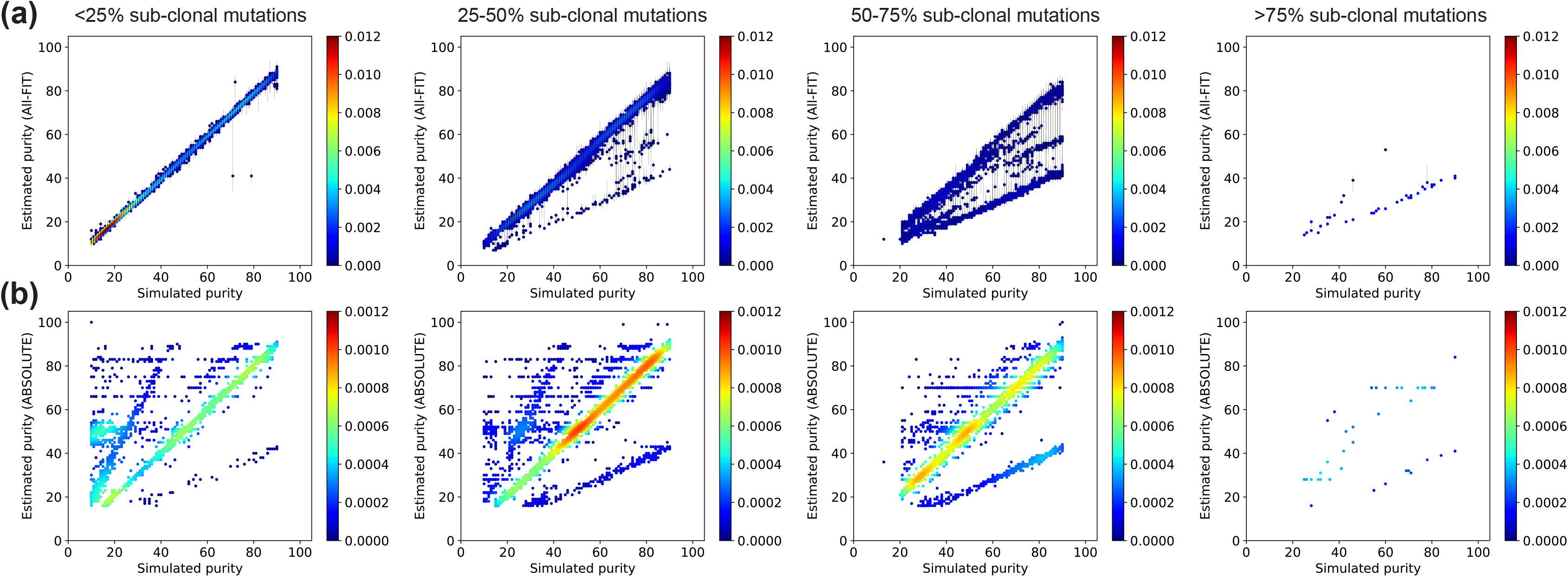
Presence of sub-clonal mutations reduces the correlation coefficient between simulated purity and estimated purity using dataset 2. (a) All-FIT accurately imputes *p* in simulated sets when percentage of sub-clonal mutations is less than 50%, beyond which, it increasingly underestimates purity at one-half of the simulated value. The correlation coefficients are respectively 0.998, 0.975, 0.764, and 0.808 from left to right panels. (b) ABSOLUTE often fails to predict *p* correctly even when the percentage of sub-clonal mutations is less than 25%. The correlation coefficients are respectively 0.745, 0.742, 0.673, and 0.562 from left to right panels. The heatmap scale represents the density of data points at each coordinate in 10,000 simulated sets.

We also tested our algorithm with a simulated dataset enriched with variants at high ploidy. Here, accuracy of All-FIT decreased to 79% while retaining the Pearson’s *r* = 0.99 (**Supplementary Figure S4a**); All-FIT performed well with nearly perfect correlation coefficient unless more than 75% of variants had ploidy > 2 with an increase in the size of the confidence intervals. Particularly, All-FIT overestimated purity at low simulated values, possibly due to relatively similar AIC weights across models, which could lead to a lack of power in breaking ambiguity (**Supplementary Figures S5a and S5c**). This could also be explained by the presence of low frequency variants that were not sub-clonal but were designated as such and excluded, leading to an increase in estimated *p*. As for ABSOLUTE, its accuracy dropped to 24% with *r* = 0.54 (**Supplementary Figure S4b**), and its correlation coefficient steadily decreased as the percentage of variants at high ploidy increased (**Supplementary Figure S5c**). It also often overestimated tumor purity regardless of the percentage of high ploidy mutations (**Supplementary Figure S5b**). ABSOLUTE’s predicted *p* with the highest likelihood mostly coupled with ploidy of two; however, the results were not significantly altered even when we removed the requirement of ploidy to be equal to two.

### Purity estimation in patient data

To test our method with patient data, we applied All-FIT to 1,861 solid tumor specimens that were sequenced using the FoundationOne assay (Foundation Medicine, Inc., Cambridge, MA). Corroborating previous studies (Carter, et al., 2012), the correlation between our computational estimates and histological values (i.e. pathological purity) was low (Pearson’s *r* = 0.28) (Figure 6). However, since all purity estimates must be interpreted in the context of tumor biology, we selected 199 tumor specimens that harbored only one hot-spot mutation in the *TP53* gene (including R175H, R248Q, R273H, R273C, R248W, R282W, R213*, G245S, Y220C, R196*, and R342*) with detected VAF > 0.10. These mutations are commonly seen in all tumor types and are known to be pathogenic. Cells with these mutations are also anticipated to have lost their wild-type copy while duplicating the mutated allele (copy-neutral LOH) (Alexandrova, et al., 2017), thus the purity of these specimens will be equal to the observed VAF of these mutations, if they are the drivers of tumor growth (Figure 1). Therefore, these driver mutations can be used as “anchors” for estimating tumor purity based on their expected biological role in cancer cells. In this analysis, we observed an improved corroboration between the estimated and anchor *p* values (*r* = 0.51). However, All-FIT overestimated purity when most variants were detected at VAF of ~0.40-0.45 (Figure 7a). This inconsistency could arise from not satisfying All-FIT’s requirement that variants from multiple mutational models should be present. For instance, in one specimen, five variants were detected, three of which had observed VAF of 0.43-0.52 and the other two were at ~0.70. All-FIT estimated tumor purity to be 0.83 (0.80-0.85) while the anchor purity from the *TP53* mutation was 0.43. If the assumption of copy-neutral LOH for this mutation was correct, this observation implied the absence of detected somatic heterozygous variants in the specimen. As expected, when we manually added a variant with VAF = 0.23, our purity estimate was corrected to 0.48 although with a large confidence interval and an unbroken ambiguity (0.45-0.66 joint with 0.81-0.83).

**Figure 6.**
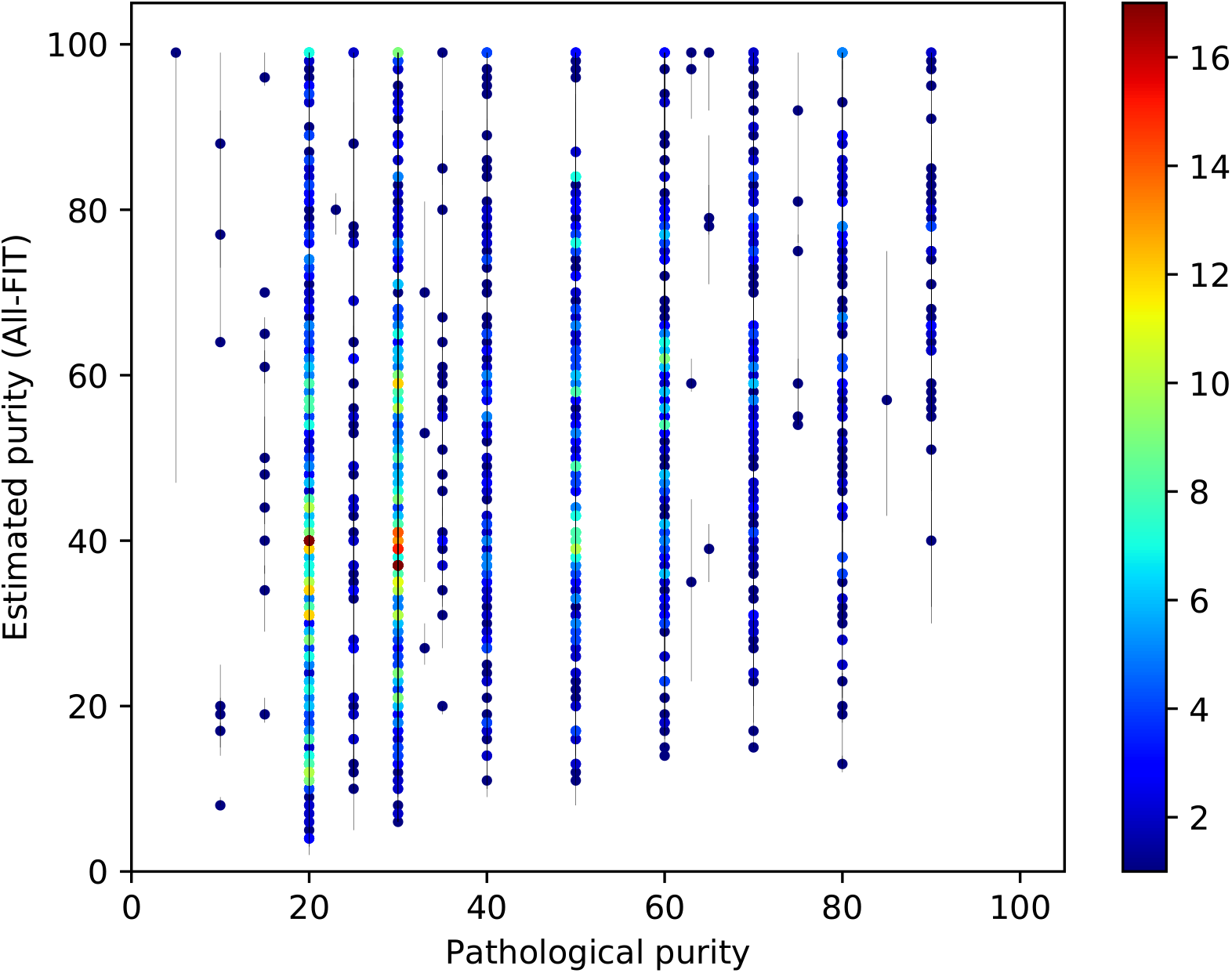
Relationship between pathological purity and estimated purity from 1,861 tumor specimens. There is limited corroboration between computational and histological estimates (*r* = 0.28). The heatmap scale represents the number of specimens at each coordinate of the graph.

**Figure 7.**
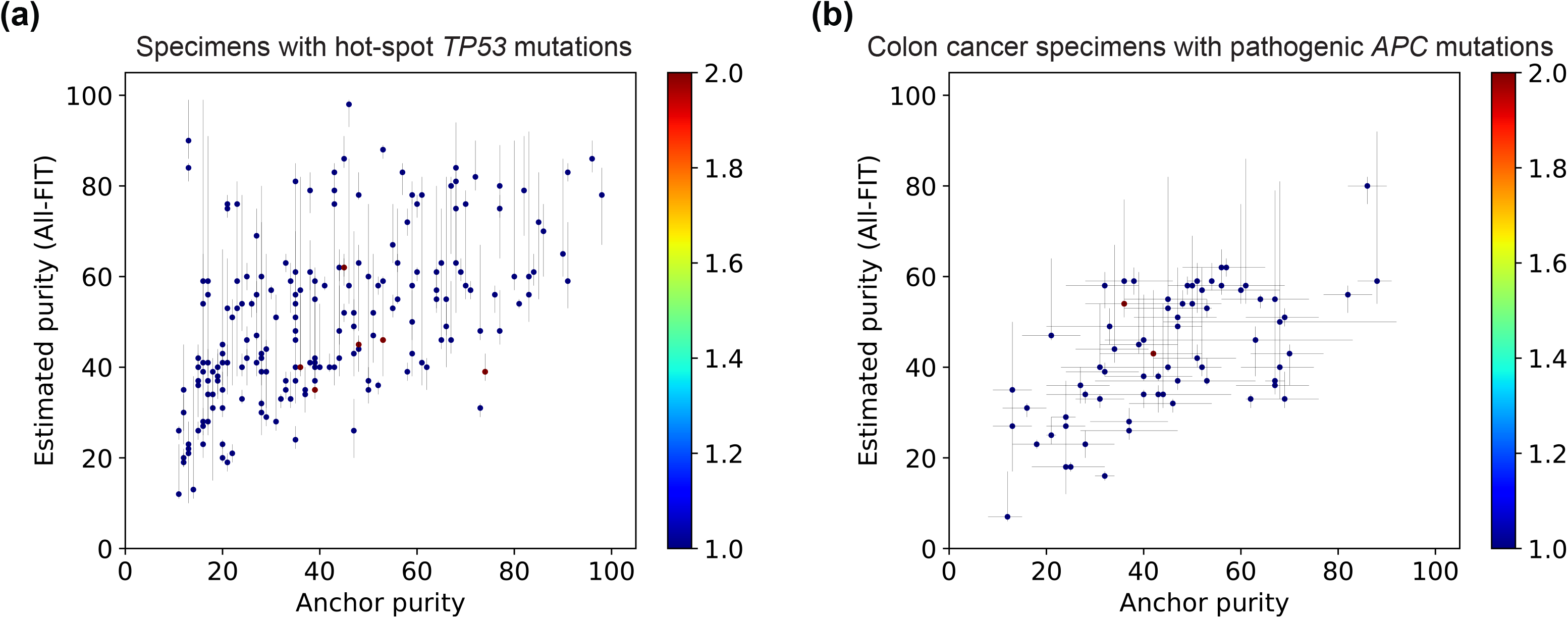
Relationship between estimated purity and allele frequency of anchor mutations. (a) All-FIT’s purity estimates corroborate with the observed VAFs of hot-spot *TP53* mutations detected in 199 solid tumor specimens, which are expected to be under copy-neutral LOH and indicative of the tumor content (*r* = 0.51). (b) All-FIT’s purity estimates corroborate with VAFs of pathogenic *APC* mutations detected in 73 colon cancer specimens (*r* = 0.60). The horizontal lines at each point represent the expected tumor content based on LOH and copy-neutral LOH models for single *APC* mutation or based on heterozygous model for two *APC* mutations. The heatmap scale represents the number of specimens at each coordinate of the graph.

In addition, as was observed in the analysis of simulated datasets, sub-clonal alterations could confound All-FIT’s results. For example, for a specimen in which the *TP53* mutation was detected with a VAF of 0.73, All-FIT estimated *p* to be 0.31 (0.29-0.32). This discrepancy was possibly due to the detection of sub-clonal variants with VAF < 0.3, which All-FIT incorrectly considered as part of the clonal population, resulting in its underestimated tumor purity. Conversely, presence of unfiltered germline variants also affected All-FIT’s estimations. This was particularly seen in a specimen in which one variant was detected at 0.07, two were detected at ~0.90, and the remaining 13 were detected at ~0.50. Although the *TP53* mutation had a VAF of 0.46, All-FIT estimated *p* to be 0.98 (0.93-0.99), as it lacked statistical power to distinguish germline and somatic variants with high confidence.

We also investigated 73 colon cancer specimens harboring pathogenic substitutions or indels in the *APC* gene. These pathogenic mutations could either undergo somatic LOH or somatic copy-neutral LOH if there is one *APC* mutation, or they could be individually somatic heterozygous if there are two *APC* mutations. There are two possible purity estimates from both one *APC* mutation and two *APC* mutations; purity with somatic LOH or somatic copy-neutral LOH with one *APC* mutation and purity based on each individual allele frequency with two *APC* mutations. In both scenarios, we considered the average of these two estimates to assess All-FIT’s results, between which we observed a correlation coefficient of 0.6 (Figure 7b). Overall, our purity estimates showed improved corroboration with anchor purity compared to those from the histological values of these specimens (**Supplementary Figure S6**).

## DISCUSSION

The complexity in cellular populations that exists within a tumor specimen is routinely summarized by the single qualitative measure of tumor purity. Since interpreting histologic as well as sequencing results relies on accurate estimates of a specimen’s tumor content, various computational solutions have been implemented to address the inaccuracies in pathological estimates (Yadav and De, 2015). These methods aim to simultaneously infer tumor purity as well as global chromosomal copy-number and often require sequencing data from pairs of tumor and normal samples. However, due to the increase use of tumor-only sequencing in precision oncology, there is a need for computational methods that can infer tumor purity of clinical specimens from detected variants, which may include somatic as well as unfiltered germline alterations. Despite the limited breadth of clinical assays that often interrogate only a few hundred genes, their high depth of sequencing often provides sufficient power to distinguish mutational models and even to infer LOH events based on variant allele frequencies.

Here, we introduced All-FIT, which offers a solution to the problem of inferring variant clonality by imputing tumor purity (*p*) from deep-sequenced specimens without matched-normal control data. It computes Akaike Information Criterion weights and cancer cell fractions for a range of somatic and germline mutational models. Through an iterative process, All-FIT estimates purity by minimizing a weighted least squared function with respect to *p* and provides statistical confidence intervals for its estimates (Figure 2a). Our application of All-FIT to patient data demonstrated the discordance between purity estimates and histological observations (Figure 6), which can be further confirmed by determining whether the mutated allele’s copy-number of each variant exceeds its own chromosomal copy-number (**Supplementary Figure S7**).

There are several caveats to our method. The first limitation is that All-FIT requires the detection of variants from various mutational models. Our analysis of simulated as well as clinical data showed that the presence of at least one somatic heterozygous variant is necessary for imputing correct tumor purity. All-FIT’s purity estimates were also greatly confounded when most detected variants were comprised of sub-clonal mutations as they violate All-FIT’s main assumption that variants are clonally present in all cancer cells. Unfiltered germline heterozygous variants also affect the statistical power to break ambiguity between multiple purity estimates, widening the confidence intervals. To address these limitations, we propose a three-step solution, where we use all detected variants to obtain a preliminary purity approximation that helps identify and exclude sub-clonal somatic as well as unfiltered germline heterozygous variants. Next, using the remaining presumably clonal events, All-FIT computes the final purity estimate and its confidence interval. All-FIT reports its results at each step to ensure completeness and to enable visual assessment by the user.

The second limitation is that we consider somatic and germline mutational models with equal probability; however, LOH and high copy-number alterations may not be detected as frequently as somatic heterozygous events. Nonetheless, All-FIT showed high accuracy and correlation with simulated ground truth even when germline variants were removed from simulated data, resulting in slightly improved accuracy and smaller confidence intervals on estimations.

The third limitation is related to the technical aspects of our method. All-FIT is restricted to model variants sequenced from hybrid-capture based assays, since interpreting VAFs from amplicon-based assays could be complicated by PCR efficiency. Furthermore, it requires specimens to have adequate admixture from surrounding normal tissues, which is at least 10%, so we simulated data with tumor purities ranging from 10% to 90%. Although All-FIT showed better performance relative to ABSOLUTE, the structure of simulated data may have worked unfavorably toward ABSOLUTE as it is not leveraged to impute tumor purity from single nucleotide variants.

## CONCLUSION

In this work, we demonstrated the robustness of a computational method for predicting tumor specimen purity without sequencing matched-normal samples. Our method is mainly applicable to clinical deep sequencing, hybrid-capture platform, which is increasingly becoming a standard approach for genomic profiling of patients in clinical settings. All-FIT can also be potentially used to estimate the abundance of tumor DNA in liquid biopsy assays. With knowledge of a specimen’s tumor content, we can now infer variant clonality and predict LOH events, leading to more tailored treatment for each patient by incorporating information on individual genes into medical decision making. Overall, our proposed method helps systematic interpretation of detected variants in a single tumor when control DNA is not available.

## FUNDING

J-WL is a pre-doctoral fellow of the New Jersey Commission on Cancer Research (DFS18PPC017). CG was a participant in the 2018 DIMACS REU program at Rutgers University supported by the National Science Foundation (CCF-1559855). HK is supported by a grant from National Cancer Institute (R01CA233662). GR, SG, and HK acknowledge support from Rutgers Cancer Institute of New Jersey (P30CA072720).

## AUTHORS’ CONTRIBUTIONS

J-WL and HK conceived the study, designed, and implanted the algorithm. J-WL and CG designed simulated datasets. FD curated patient clinical data. GR and SG performed clinical interpretation of the results. HK supervised the study. All authors contributed to the drafting of the manuscript and critical discussion of the results. All authors read and approved the final manuscript.

## CONFLICT OF INTEREST

GR serves on a scientific advisory board and as consultant to Personal Genome Diagnostics. SG serves on a scientific advisory board and as consultant for Inspirata Inc, holds patents on digital imaging technology licensed to Inspirata Inc, holds equity in Inspirata Inc, serves on an advisory board for Novartis Pharmaceuticals, and serves as a consultant for Roche and Foghorn Therapeutics. Other authors declare that they have no conflict of interests.

## ACKNOWLEDGMENTS

The authors would like to thank Nahed Jalloul and other members of the Khiabanian Lab and the staff, physicians, and pathologists of the Division of Precision Medicine Oncology at Rutgers Cancer Institute of New Jersey.

---

**Supplementary Figure S1.** The effect of total number of mutations in a sample on the accuracy of All-FIT, using dataset with similar criteria as dataset 1, but with the number of variants in each set ranging from 5 to 100. The x-axis represents the number of mutations in interval of 5, and the y-axis represents the difference between estimated *p* and simulated *p*, without considering the confidence intervals. The accuracy of All-FIT is independent of the number of mutations, when there are at least 15 variants per sample.

**Supplementary Figure S2.** Relationship between simulated purity (ground truth) and estimated purity from All-FIT using dataset 1, after eliminating germline variations from the simulated sets. All-FIT’s accuracy improves to 92% with correlation coefficient of 0.99. The heatmap scale represents the density of data points at each coordinate of the graph from 10,000 simulated sets.

**Supplementary Figure S3.** Presence of sub-clonal mutations reduces the correlation coefficient between simulated purity and estimated purity using dataset 2, with only somatic variants present. As shown in Figure 5, All-FIT accurately imputes *p* in simulated sets when percentage of sub-clonal mutations is less than 50%, beyond which, an increasing number of simulated sets is underestimated at one-half of simulated values. However, having matched-normal control provides a better predictive power to All-FIT by increasing the overall correlation coefficient and less simulated sets are predicted with incorrect *p*. The correlation coefficients are respectively 0.999, 0.985, 0.837, and 0.719 from upper left to lower right panels.

**Supplementary Figure S4.** Relationship between simulated purity (ground truth) and estimated purity using dataset 3. (a) All-FIT shows almost perfect correlation (*r* = 0.99) with an accuracy of 79%. All-FIT overestimates for some simulated sets at low end of purity; this can be explained by the misclassification of low frequency variants as sub-clonal mutations. (b) ABSOLUTE exhibits a lower correlation of *r* = 0.54 with 24% accuracy. Most simulated sets are imputed with incorrect *p*, which is either two times or one-half of the simulated purity.

**Supplementary Figure S5.** Presence of high ploidy mutations has minimal impact on the correlation coefficient between simulated purity and estimated purity from All-FIT. (a) Overall, aneuploidy has no significant effect on All-FIT, apart from some simulated sets being estimated at high *p* at low end of simulated purity spectrum when >75% of variants have high ploidy. (b) ABSOLUTE overestimates tumor purity regardless of the percentage of high ploidy variants, and more simulated sets are underestimated when the percentage is higher than 50%. (c) All-FIT shows nearly perfect correlation when <80% of variants have high ploidy, but ABSOLUTE’s correlation steadily decreases from 0.79, due to its strong dependence on copy-number change instead of single variant change.

**Supplementary Figure S6.** Correlation between estimated purity and pathological purity for (a) solid tumor specimens with hot-spot *TP53* mutations (*r* = 0.22) and (b) colon cancer specimens with pathogenic *APC* mutations (*r* = 0.32). The heatmap scale represents the number of specimens at each coordinate of the graph.

**Supplementary Figure S7.** Discordance between purity estimates and histology in a clinical sample. The x-axis represents chromosomal copy-number or ploidy (*Y*) and the y-axis represents the mutated allele’s copy-number (*c*_m_) calculated based on its observed VAF, *Y*, and *p* (Figure 1), with (a) using estimated *p* = 0.75 as purity and (b) using histological value of 0.3 as purity. At *p* = 0.75, most variants have *c*_m_ (in integer) ≤ *Y*, which is further justified by the *TP53* mutation detected at VAF of 0.77.

## REFERENCES

Alexandrova, E.M., et al. p53 loss-of-heterozygosity is a necessary prerequisite for mutant p53 stabilization and gain-of-function in vivo. Cell Death Dis 2017;8(3):e2661.

Amirouchene-Angelozzi, N., Swanton, C. and Bardelli, A. Tumor Evolution as a Therapeutic Target. Cancer Discov 2017.

Andor, N., et al. EXPANDS: expanding ploidy and allele frequency on nested subpopulations. Bioinformatics 2014;30(1):50–60.

Boeva, V., et al. Control-FREEC: a tool for assessing copy number and allelic content using next-generation sequencing data. Bioinformatics 2012;28(3):423–425.

Brok, W.D.d., et al. Homologous Recombination Deficiency in Breast Cancer: A Clinical Review. JCO Precision Oncology 2017(1):1–13.

Carter, S.L., et al. Absolute quantification of somatic DNA alterations in human cancer. Nat Biotechnol 2012;30(5):413–421.

Damodaran, S., Berger, M.F. and Roychowdhury, S. Clinical tumor sequencing: opportunities and challenges for precision cancer medicine. Am Soc Clin Oncol Educ Book 2015:e175–182.

Foran, D.J., et al. Roadmap to a Comprehensive Clinical Data Warehouse for Precision Medicine Applications in Oncology. Cancer Inform 2017;16:1176935117694349.

Frampton, G.M., et al. Development and validation of a clinical cancer genomic profiling test based on massively parallel DNA sequencing. Nat Biotechnol 2013;31(11):1023–1031.

Garraway, L.A. Genomics-driven oncology: framework for an emerging paradigm. J Clin Oncol 2013;31(15):1806–1814.

Gusnanto, A., et al. Correcting for cancer genome size and tumour cell content enables better estimation of copy number alterations from next-generation sequence data. Bioinformatics 2012;28(1):40–47.

Khiabanian, H., et al. Inference of Germline Mutational Status and Evaluation of Loss of Heterozygosity in High-Depth, Tumor-Only Sequencing Data. JCO Precis Oncol 2018;2018.

Li, M.M., et al. Standards and Guidelines for the Interpretation and Reporting of Sequence Variants in Cancer: A Joint Consensus Recommendation of the Association for Molecular Pathology, American Society of Clinical Oncology, and College of American Pathologists. J Mol Diagn 2017;19(1):4–23.

McGranahan, N., et al. Allele-Specific HLA Loss and Immune Escape in Lung Cancer Evolution. Cell 2017;171(6):1259–1271 e1211.

Pawlyn, C., et al. Loss of heterozygosity as a marker of homologous repair deficiency in multiple myeloma: a role for PARP inhibition? Leukemia 2018.

Ptashkin, R.N., et al. Prevalence of Clonal Hematopoiesis Mutations in Tumor-Only Clinical Genomic Profiling of Solid Tumors. JAMA Oncol 2018.

Riedlinger, G., et al. Association of JAK2-V617F Mutations Detected by Solid Tumor Sequencing With Coexistent Myeloproliferative Neoplasms. JAMA Oncol 2019.

Roth, A., et al. PyClone: statistical inference of clonal population structure in cancer. Nat Methods 2014;11(4):396–398.

Sade-Feldman, M., et al. Resistance to checkpoint blockade therapy through inactivation of antigen presentation. Nat Commun 2017;8(1):1136.

Severson, E.A., et al. Detection of clonal hematopoiesis of indeterminate potential in clinical sequencing of solid tumor specimens. Blood 2018;131(22):2501–2505.

Shaw, K.R.M. and Maitra, A. The Status and Impact of Clinical Tumor Genome Sequencing. Annu Rev Genomics Hum Genet 2019.

United States Food & Drug Administration. FDA announces approval, CMS proposes coverage of first breakthrough-designated test to detect extensive number of cancer biomarkers. In. www.fda.gov; 2017.

Van Loo, P., et al. Allele-specific copy number analysis of tumors. Proc Natl Acad Sci U S A 2010;107(39):16910–16915.

Yadav, V.K. and De, S. An assessment of computational methods for estimating purity and clonality using genomic data derived from heterogeneous tumor tissue samples. Brief Bioinform 2015;16(2):232–241.

