## Supplementary figures and images for "All-FIT: Allele-Frequency-based Imputation of Tumor Purity from High-Depth Sequencing Data"

### Supplementary Figures

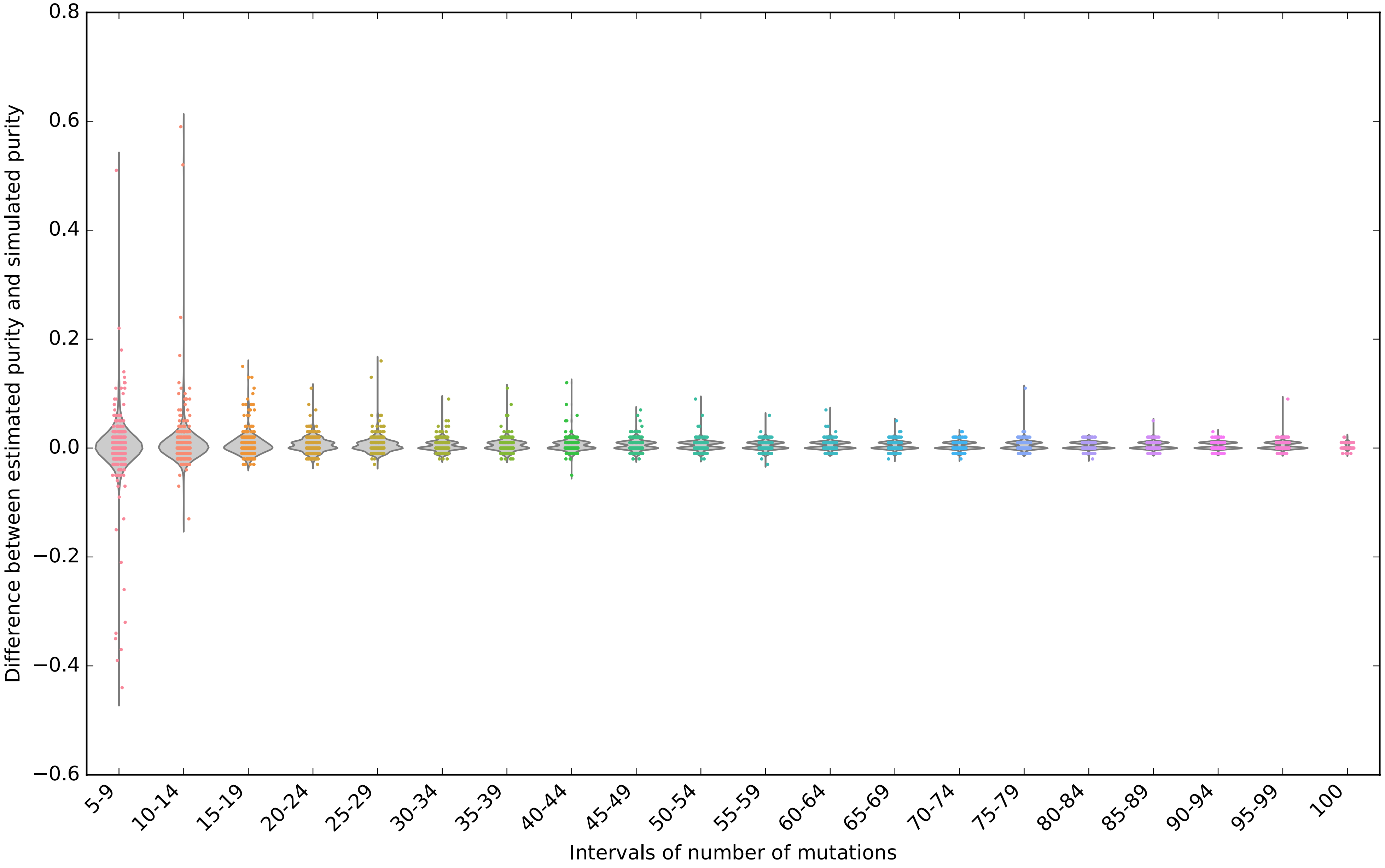

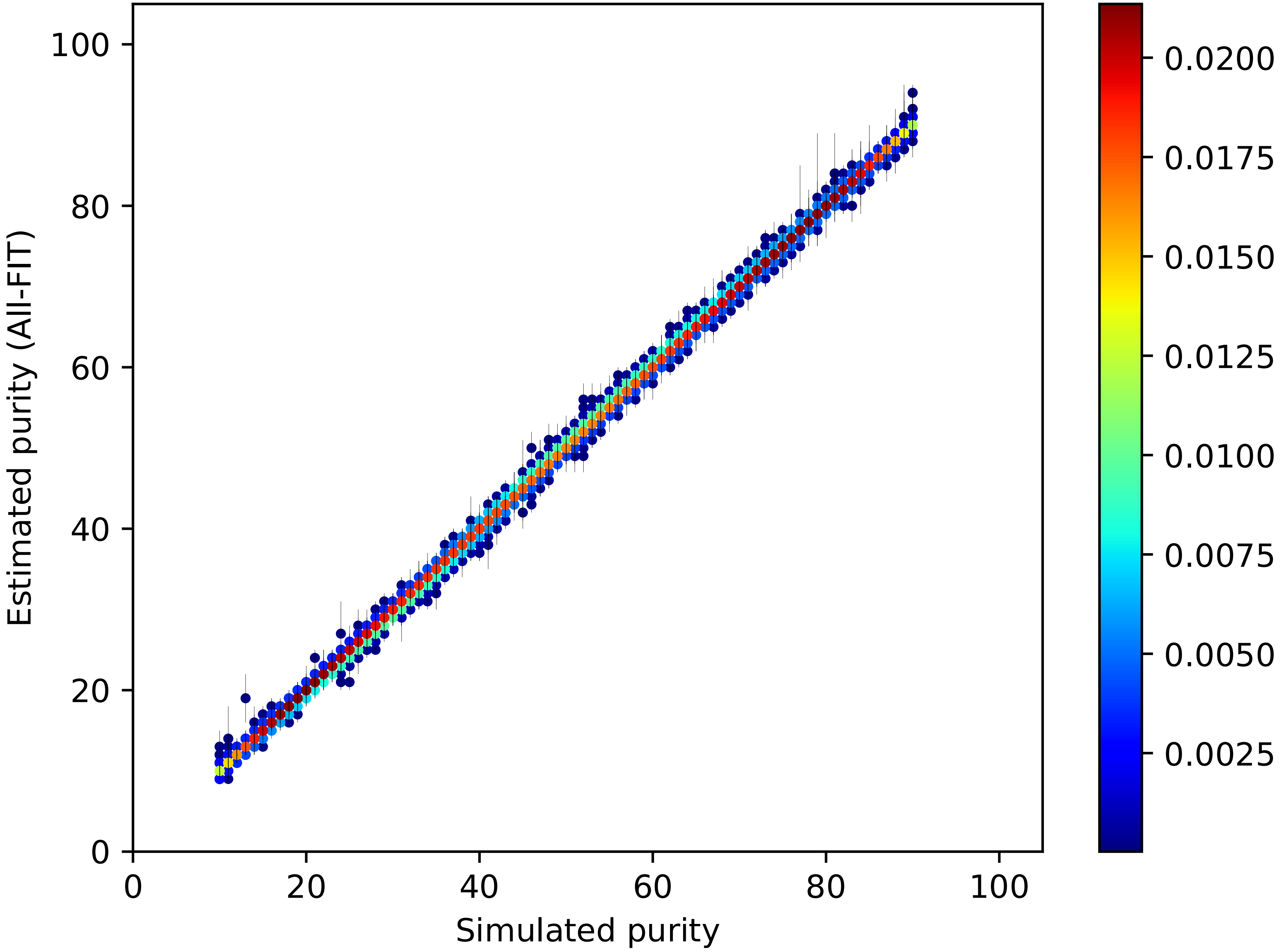

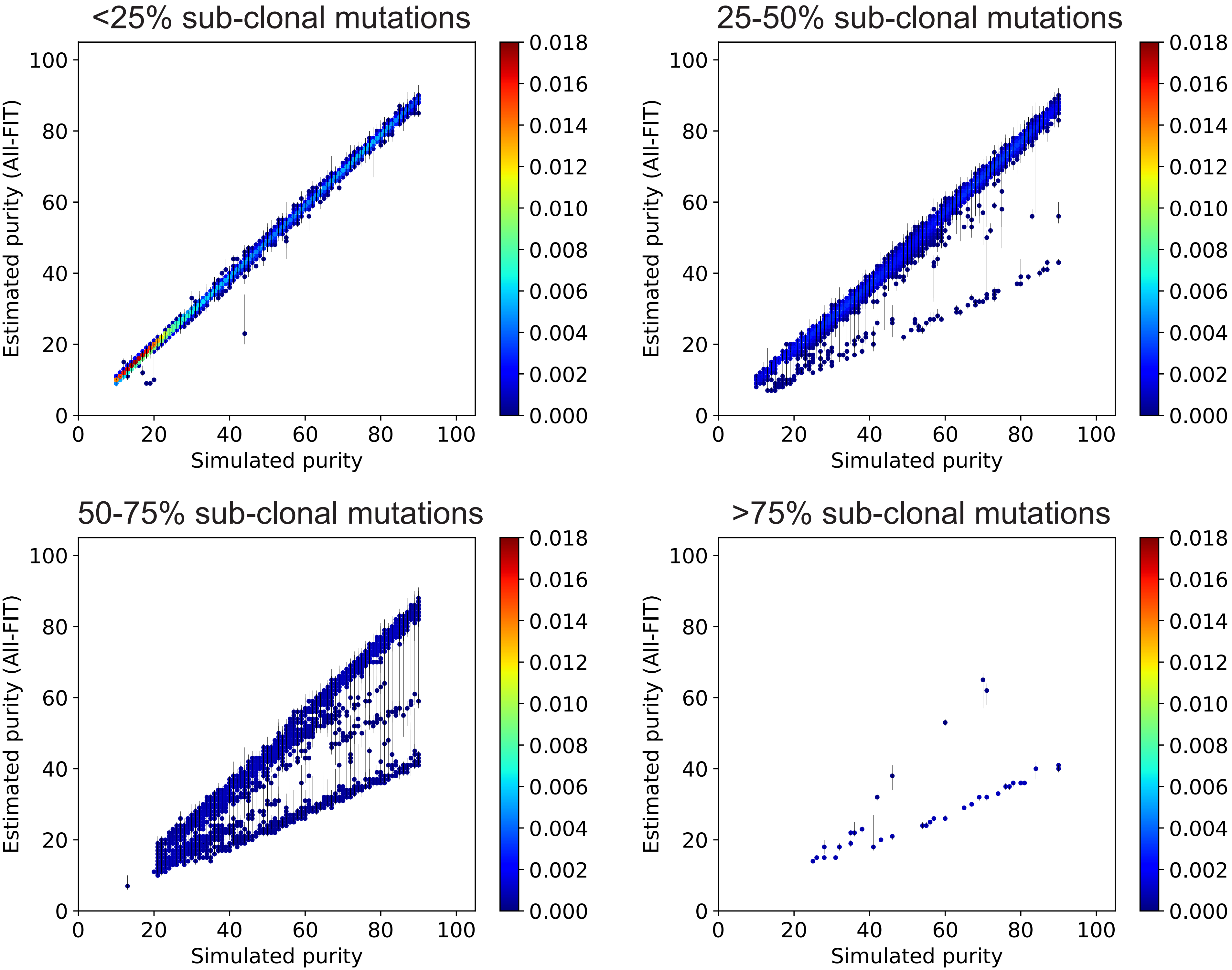

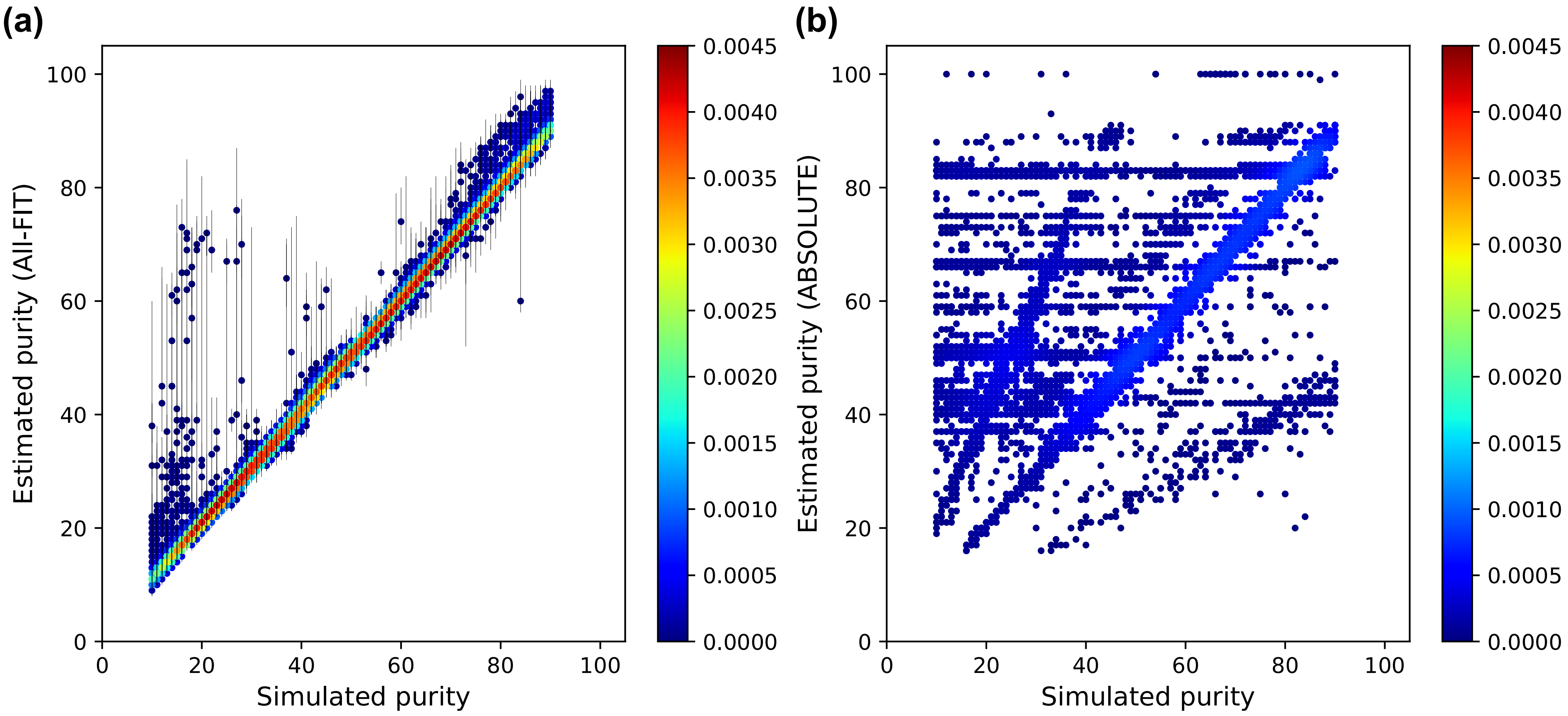

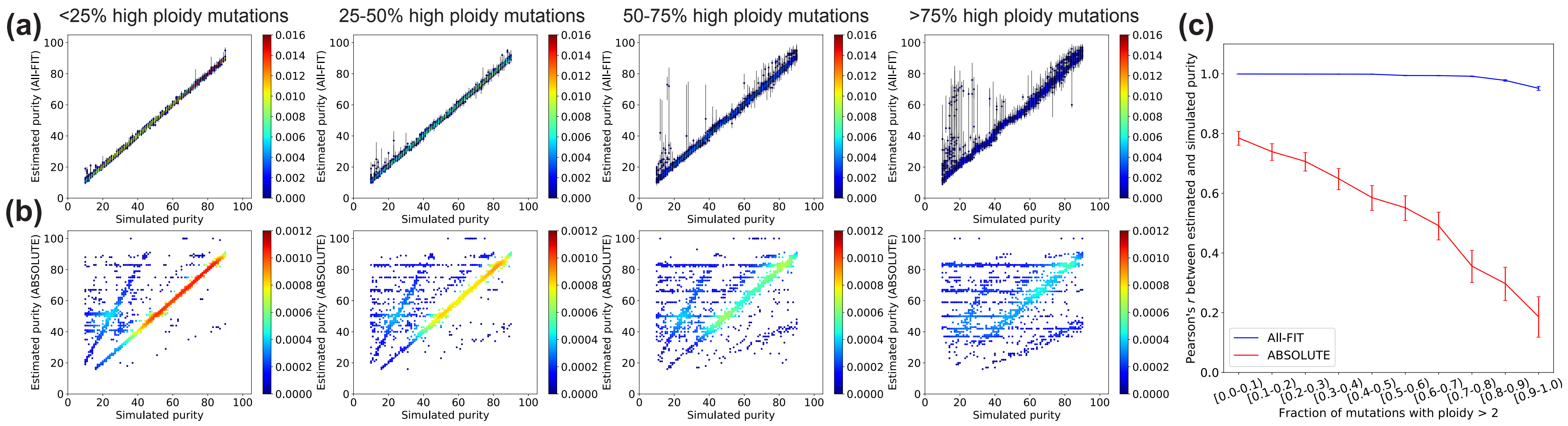

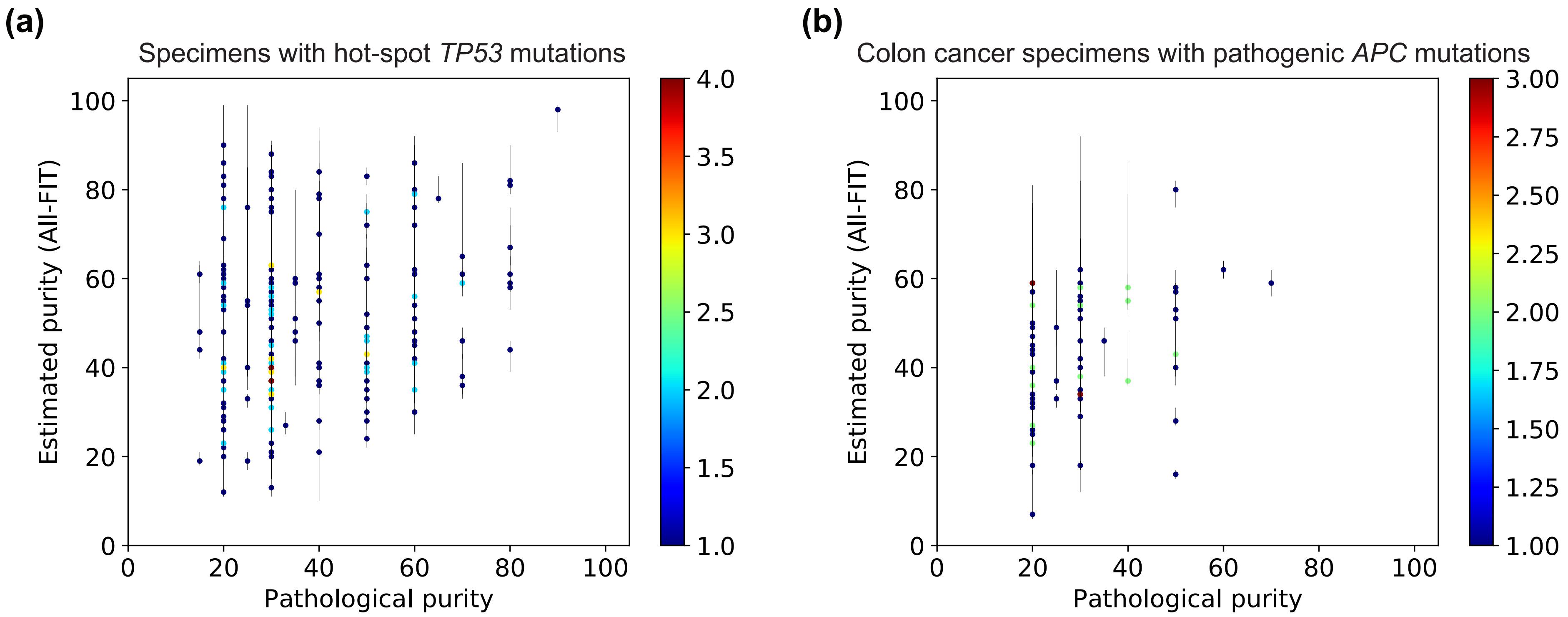

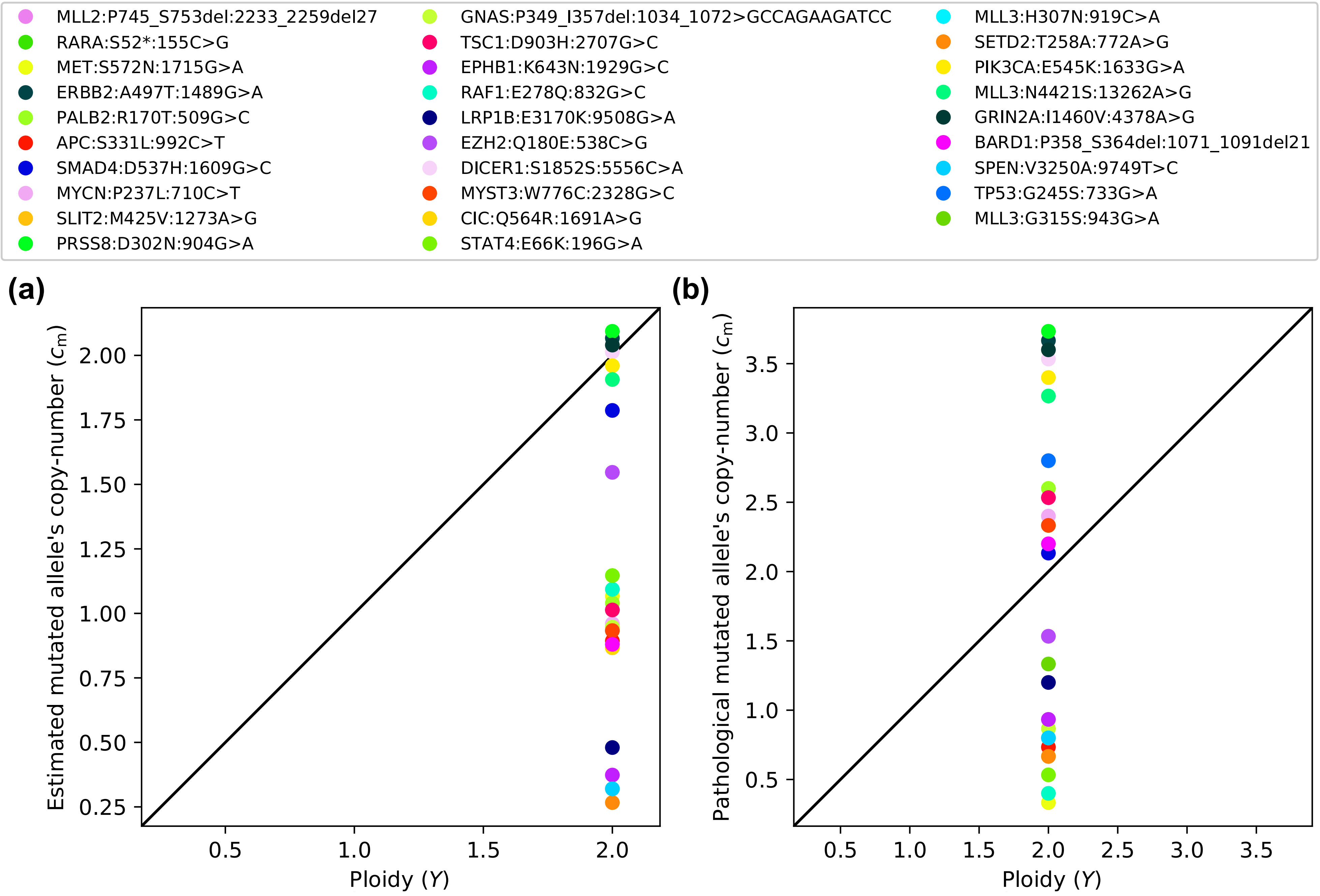
